# Evaluation of a Loop-Mediated Isothermal Amplification assay to detect carbapenemases from *Acinetobacter* spp. directly from bronchoalveolar lavage fluid

**DOI:** 10.1101/2020.05.11.089904

**Authors:** J. Moreno-Morales, A. Vergara, T. Kostyanev, J. Rodriguez-Baño, H. Goossens, J. Vila

## Abstract

Carbapenem-resistant *Acinetobacter* spp. mainly *Acinetobacter baumannii* are frequently causing nosocomial infections with high mortality. In this study, the efficacy of the Eazyplex^®^ SuperBug Complete A system (Amplex Diagnostics GmbH, Gars-Bahnhof, Germany), based on loop-mediated isothermal amplification (LAMP), to detect the presence of carbapenemases in *Acinetobacter* spp. directly from bronchoalveolar lavage samples was assessed, detecting all tested carbapenemases in less than 30 minutes with a sensitivity of 10^3^ CFU/ml.

## Main body

Carbapenems are potent β-lactam antibiotics with broad-spectrum and bactericidal mode of action (1). Their use was increased due to the spread of extended spectrum β- lactamase producing Enterobacteriaceae towards whom they are active (2, 3). Carbapenems are considered one of the most efficacious antimicrobials to treat bacterial infections (1). However, resistance by carbapenemases did not take long to appear and it poses a major threat to public health. (2)

*Acinetobacter* spp. members and specially carbapenem-resistant *Acinetobacter baumannii* (CRAB) are among the world’s most dangerous pathogen threats. CRAB has been classified as a critical priority pathogen by the WHO’s priority pathogens to guide R&D of new antibiotics (4, 5) and as an urgent threat that requires aggressive action by CDC (6).

Even though community-acquired *Acinetobacter* infections can occur, the most common and acute infections happen in the nosocomial setting. *Acinetobacter* lurks around intensive care units and surgical wards causing a number of infections (e.g., on burns and soft tissue, urinary tract, bloodstream, etc.) and specially ventilator-associated pneumonia (VAP) in patients under mechanical ventilation (3, 7)

VAP develops in intensive care units in patients under ventilation for at least 48h. Rapid diagnostic of VAP causing pathogens is of utmost importance: VAP patients not only have longer hospital stays and need more antibiotics, therefore their treatment is more expensive, but also have higher mortality (8, 9).

This study aims to evaluate Eazyplex^®^ Superbug Complete A based on loop-mediated isothermal amplification (LAMP) for detecting carbapenemase produced by *Acinetobacter* directly from inoculated BAL samples.

## Materials and Methods

Eazyplex^®^ SuperBug Complete A kit, used with the Genie II device, is a molecular diagnostics kit that detects a selection of genes which express carbapenemases (including metallo-β-lactamases and oxacillinases). Results after detection of bacterial DNA are presented within 30 minutes.

The kit is composed of 8 tube strips each with a mix of lyophilized agents for the amplification of one of the following 7 genes: *bla*_KPC_, *bla*_NDM_, *bla*_VIM_, *bla*_OXA-48_, *bla*_OXA-23_, *bla*_OXA-40_ and *bla*_OXA-58_. The eighth tube being an internal inhibitory control.

Once the samples are prepared and the strip is inside the Genie II device, a LAMP is performed. The reaction is incubated at 66 C º for 30 minutes and detection is performed via fluorescence excitation, for up to two strips at a time.

A total of 11 *Acinetobacter* spp. strains producing OXA-23, OXA-40, OXA-58 and NDM were selected (Table). Isolate identification was performed via MALDI-TOF/MS (Bruker Daltonics, Bremen, Germany). Carbapenemase gene detection was checked via conventional PCR for each of the strains (10, 11).

Negative BAL samples were identified and collected at the Clinical Microbiology Laboratory from Hospital Clinic of Barcelona; samples were stored at – 80 C °. McFarland solutions were prepared from each of the 11 *Acinetobacter* strains and serial dilutions in saline solution were made to finally spike BAL samples to a concentration of 10^2^ and 10^3^ CFU/ml.

The protocol consisted in: centrifugation of 850 μL of the 10^2^ and 10^3^ spiked BAL samples (at 14,000 g for 5 minutes), addition of 500 μL of resuspension and lysis fluid (RALF, provided with the kit) to the pellet obtained, incubation at 99°C for 2 minutes and a final centrifugation step (4,000 rpm for 2 minutes). Finally, 25 μL of the supernatant were added to each tube of the assay strip. The hands-on time took a maximum of 15 minutes per strain.

## Results and discussion

Increasing resistance to antimicrobials and specifically carbapenems is reported in *A. baumannii* in past years. In Spain, there has been an increase of up to 40% in *A. baumannii* clinical isolates presenting resistance to carbapenems from 2000 to 2010; 86% of the 446 *A. baumannii* clinical isolates presented resistance to carbapenems in 2010’s study (12, 13).

This situation is rather common, recently ECDC has reported ≥50% of *Acinetobacter* spp. invasive isolates present resistance to carbapenems in Hungary, Poland, Bulgaria, Latvia, Italy, Spain, Cyprus, Romania, Lithuania, Greece and Croatia in 2018’s Annual report of the European Antimicrobial Resistance Surveillance Network (14).

Current effective antibiotics for the treatment of CRAB are scarce and are not the most suitable therapeutic agents due to poor pharmacokinetics, toxicity (as in the case of polymyxins) and emergence of resistance (15, 16). Chromosome and/or plasmid encoded carbapenemases are the main mechanism of resistance to carbapenems in CRAB (3, 17), thus rapid detection of carbapenemases is key to guide effective antibiotic therapies (16). Detection of carbapenemase producing genes in the tested strains using Eazyplex^®^ SuperBug Complete A assay is shown in the Table and the results agree with conventional PCR results. Detection time values vary per strain and gene. Only 5 concentrations tested out of 22 did not provide detection of the carbapenemase producing gene, all being at the lowest concentration tested at 10^2^ CFU/ml, therefore, the limit of sensitivity is 10^3^ CFU/ml.

With a maximum hands-on time of 15 minutes per sample and 30 minutes run time (approximately 45 minutes total), this assay proves to be a great advantage compared to routine methods in the clinical microbiology laboratory that need 16-24h for results to be obtained. Naturally, further antimicrobial susceptibility testing should be considered in all samples.

We visualize the following workflow for diagnosis of hospital-acquired pneumonia (HAP): when the sample arrives to the clinical microbiology laboratory, rapid identification of the bacteria causing HAP is performed also using a LAMP reaction approach (18), if *A. baumannii* is identified as the pathogen causing the infection, the method to detect specific carbapenemases in *Acinetobacter* described in this study is performed.

## Conclusion

Using Eazyplex^®^ SuperBug Complete A assay will allow to guide and optimize antibiotic therapies earlier than with usual techniques used in the laboratory, which it likely mean a decrease in mortality. This assay represents the kind of advantages that investing in molecular diagnostics brings to the clinical practice: allows the identification of specific resistance mechanisms in approximately 45 minutes and if sample identification using LAMP was included as a first step (1h), both pathogen and resistance mechanism could be identified in less than 2h.

## Acknowledgements

We acknowledge support from the Spanish Ministry of Science and Innovation through the “Centro de Excelencia Severo Ochoa 2019-2023” Program (CEX2018-000806-S), and support from the Generalitat de Catalunya through the CERCA Program. Supported by Plan Nacional de I+D+i 2013-2016 and Instituto de Salud Carlos III, Subdirección General de Redes y Centros de Investigación Cooperativa, Ministerio de Economía, Industria y Competitividad, Spanish Network for Research in Infectious Diseases (REIPI RD16/0016/0010) - co-financed by European Development Regional Fund “A way to achieve Europe”, Operative program Intelligent Growth 2014-2020. The research leading to these results was conducted as part of the COMBACTE-CARE project.

## Transparecy declarations

Nothing to disclose.

---

| Strain | Resistance gene (*bla*) | Concentration (CFU/ml) | Eazyplex® Superbug Complete A (minutes:seconds) |
| --- | --- | --- | --- |
| *A. baumannii* 5075 | OXA-23 | 10<sup>3</sup> | 14:00 |
|  |  | 10<sup>2</sup> | 16:15 |
| *A. baumannii* R308 | OXA-23 | 10<sup>3</sup> | 11:45 |
|  |  | 10<sup>2</sup> | - |
| *A. baumannii* 2a1 | OXA-23 | 10<sup>3</sup> | 28:00 |
|  |  | 10<sup>2</sup> | 21:15 |
| *A. baumannii* R180 | OXA-40 | 10<sup>3</sup> | 12:50 |
|  |  | 10<sup>2</sup> | 13:15 |
| *A. baumannii* R329 | OXA-40 | 10<sup>3</sup> | 13:30 |
|  |  | 10<sup>2</sup> | - |
| *A. baumannii* 67 | OXA-40 | 10<sup>3</sup> | 11:30 |
|  |  | 10<sup>2</sup> | 17:00 |
| *A. baumannii* R247 | OXA-58 | 10<sup>3</sup> | 16:10 |
|  |  | 10<sup>2</sup> | 18:30 |
| *A. baumannii* R80 | OXA-58 | 10<sup>3</sup> | 16:15 |
|  |  | 10<sup>2</sup> | - |
| *A. baumannii* 81 | OXA-58 | 10<sup>3</sup> | 16:45 |
|  |  | 10<sup>2</sup> | 21:30 |
| *Acinetobacter pittii* M2 | NDM* | 10<sup>3</sup> | 20:00 |
|  |  | 10<sup>2</sup> | - |
| *Acinetobacter dijkshoorniae* M1 | NDM* | 10<sup>3</sup> | 18:30 |
|  |  | 10<sup>2</sup> | - |

Eazyplex® Superbug Complete A detection time per concentration tested for each of the *Acinetobacter* spp. strains tested. ‘-’ indicates no detection of the carbapenemase producing gene in the concentration tested.

